# Absence-like epileptic spike-wave discharges in adult Lister Hooded rats and their pharmacological modulation

**DOI:** 10.1101/2020.12.03.409433

**Authors:** Gabriella Nyitrai, Pálma Diószegi, Gergely Somogyi, András Czurkó

## Abstract

Efforts to advance translation through pre-clinical behavioural and pharmacological tests prompted attention to rat strain differences. Particularly the use of touchscreen technology for cognitive testing initiated the widespread use of Lister Hooded and Long Evans rats and they differed in pharmacological sensitivity to certain drugs. One possible reason for this rat strain difference could be that Long Evans rats produce high-amplitude spike-wave discharges (SWDs) in their cortical EEG recordings, while no information available about Lister Hooded rats in this regard. As a serendipitous observation, we noticed the presence of SWDs during the EEG recordings of Lister Hooded rats. In this study, therefore, we examined these spontaneous SWDs in two groups of Lister Hooded rats. The number and sum duration of the SWDs were similar to that was observed in other rat strains. We found SWDs during wakefulness, slow-wave sleep (SWS) and rapid eye movement (REM) sleep, their duration was the longest during wakefulness, but their number and sum duration were also high during REM. The GABA-B receptor agonist baclofen exacerbated, while the GABA-B antagonist SCH50911 reduced the occurrence of the recorded SWDs. Typical anti-seizure medications, valproate and diazepam, decreased the number and sum duration of SWDs. Although the two rat strains typically used in touchscreen experiments are similar in term of SWDs, the occurrence and possible pharmacological modulation of SWDs are considerable during their use in behavioural experiments.

## 1. Introduction

Efforts to improve pharmaceutical discovery for the identification of effective treatments for cognitive dysfunctions and to advance translation from pre-clinical to clinical research has prompted attention to rat strain variations in behavioural and pharmacological tests (Kumar et al., 2015; McDermott and Kelly, 2008; Mohler et al., 2015; Stensbol and Kapur, 2015). Particularly the use of the touchscreen technology initiated the widespread use of Lister Hooded and Long Evans rats in behavioural pharmacology studies as their visual acuity is better than those of albino rats (Kumar et al., 2015). On the other hand, Lister Hooded rats seem to be more sensitive than Long–Evans rats to drugs traditionally used to cause cognitive impairments (Mohler et al., 2015).

One possible reason for this rat strain difference is that Long Evans rats produce high-amplitude rhythmical cortical EEG waves with spikes, also called polyspike plus spike-and-wave discharges, during awake immobility (Kaplan, 1985; Semba et al., 1980; Vanderwolf, 1975) while no information available about this EEG activity in Lister Hooded rats.

This particular EEG activity in Long Evans rats, also known as paroxysmal 7–12 Hz high-voltage rhythmic spike discharge and often considered to be the EEG correlate of absence-like seizure activity that resembles the human petit mal epilepsy (Kaplan, 1985; Polack and Charpier, 2006; Shaw, 2004, 2007; van Luijtelaar et al., 2011). This absence-like seizure activity in the EEG of the rat denotes synchronized spontaneously occurring rhythmic thalamocortical oscillations characterized by spike-and-wave discharges (SWDs) and have been described in various rat strains (Buzsaki et al., 1988b; Jando et al., 1995; Shaw, 2004; Willoughby and Mackenzie, 1992). These discharges have relatively high EEG amplitude (300-1000 μV), a dominant frequency in the theta-alpha range (6-12 Hz) and the duration from 1 sec to several seconds (Buzsaki et al., 1988a; Jando et al., 1995). The animals stay inactive during these discharges, similarly to the human absence episodes (Depaulis and Charpier, 2018; Panayiotopoulos, 1999).

SWDs differ in their sum duration and number widely among rat strains as having been demonstrated in various EEG studies (Jando et al., 1995; Shaw, 2004; Willoughby and Mackenzie, 1992). Moreover, two rat strains with robust spontaneous SWD activity were selected and used in the preclinical research as animal models of the human absence epilepsy, the GAERS (Danober et al., 1998; Marescaux et al., 1992b; Vergnes et al., 1982) and the Wag/Rij (Coenen and Van Luijtelaar, 2003; Drinkenburg et al., 1991) rat strains. To some extent, Long-Evans rats are also well-characterized and validated as a model for absence epilepsy (Kozak et al., 2020; van Luijtelaar et al., 2011), although thousands of them have been subjects of psychological experiments throughout the decades (Kaplan, 1985). Consequently from early on, others have proposed that these SWD oscillations in the Long Evans rat are analogous to human mu rhythms and they represent physiological activity that does not preclude perception (Wiest and Nicolelis, 2003).

The Lister Hooded rat strain is one of the most widely used rat strain in behavioural pharmacology studies (McDermott and Kelly, 2008; Mohler et al., 2015), and to our knowledge, no SWD oscillations described in the EEG recordings of this rat strain. Measurement of the rodent EEG activity offers adequate translational power, as EEG oscillations are analogous in rodents and humans during the sleep stages and show similar changes following drug administration (Babiloni et al., 2013; Drinkenburg et al., 2015). As a serendipitous observation, we noticed the presence of SWDs during the EEG recordings of Lister Hooded rats. As a characterization of the basal EEG activity of Lister-Hooded rats is missing from the available literature, we decided to quantify these SWDs and characterize their pharmacological modulation.

Absence epilepsy in rat models has a similar pharmacological profile as humans (Manning et al., 2003; Vrielynck, 2013). Drugs enhancing GABAergic inhibition could intensify SWDs, while drugs that are antagonists at the GABA-B receptor decreasing these pathological oscillations (Marescaux et al., 1992a; Ritchie et al., 2003; van Luijtelaar et al., 2002; Vergnes et al., 1997). Diazepam could also decrease SWDs in animals with a less known mechanism (Micheletti et al., 1985; van Luijtelaar et al., 2002), while valproate is a widely accepted and effective anti-seizure medication in both animals and humans (Manning et al., 2003; Marescaux et al., 1985; Vrielynck, 2013).

Therefore, we evaluated the EEG of Lister Hooded rats focusing the occurrence of probable SWDs and explored the effects of drugs known to modulate SWDs, as an agonist and an antagonist of the GABA-B receptor as well as anti-seizure medications, diazepam and valproate.

## 2. Methods

### 2.1. Drugs

*R,S*-baclofen (Tocris), SCH50911 (Tocris), sodium-valproate (synthetized by Gedeon Richter Plc., batch number: MKCD3350) and diazepam (HCl salt, synthetized by Gedeon Richter Plc., batch number: L77028F), was dissolved in physiological saline. All solutions were prepared freshly, 1 hour before application.

### 2.2. Animals and surgery

Male Lister Hooded rats (n=17=11+6, WOBE/Envigo, , weights 270-330 gr, 8-9 week-old) were implanted with stainless steel screw electrodes (1.4×4.0 mm) over the right frontal (A: 2.0 mm, L: 2.0 mm) left parietal (A: −3.0 mm, L: 2.0 mm) left occipital (A: −7.0 mm, L: 1.5 mm) and right visual (A: −5.5 mm, L: 5.0 mm) cortices under isoflurane anesthesia (1.5-2.0% in air). Coordinates refer mm relative to bregma according to the Rat Brain Atlas of Paxinos & Watson (Paxinos G, 2005). A reference and a separate ground electrode were implanted over the cerebellum. To monitor EMG activity, an insulated stainless steel wire was inserted into the neck nuchal muscle. Electrodes were soldered to a connector and secured to the skull with dental cement. Rats received 0.2 ml Baytril (0.5 mg/kg, KVP, Pharma- and VeterianerProdukte GmbH, Kiel) and 0.2 ml Norocarp (0.5 mg/ml, Norbrook Lab. LTD, Station Works) injections *s.c*.at the end of the surgery and on the following two days. After a surgical recovery of a minimum of 2 weeks, animals were acclimatized to the handling procedures.

Animals are kept under standard, temperature and humidity controlled laboratory conditions with food and water ad libitum. Throughout the whole experiment, animals were kept in a 12h light/dark cycle, with the light on at 6:00 a.m.

Experiments were carried out in accordance with the European Communities Council Directive of 24 November 1986 (86/609/EEC) and were carried out in strict compliance with the European Directive 2010/63/EU regarding the care and use of laboratory animals for experimental procedures. All the procedures conformed to the guidelines of the National Institutes of Health and the guidelines of the local Animal Care and Use Committee and were approved by the local Ethical Committee of Gedeon Richter Plc. All efforts were made to minimize the number of animals as well as pain or discomfort.

### 2.3. EEG recordings

Rats were placed into individual recording cages (same size as their home cage) and connected to the amplifiers (through swivels) with flat and flexible connecting cables allowing rats to continue their regular activity. Recording sessions started at 9:00 a.m. and lasted for 4 hours.

EEG signals were filtered (0.16-1000 Hz), amplified (5000x) with 8-channel EEG amplifiers (Supertech Ltd., Pécs, Hungary), digitalized with CED 1401 data acquisition system at a rate of 2 kHz (Cambridge Electronic Design Ltd., UK) and recorded with Spike2 (version 7.4, Cambridge Electronic Design Ltd., UK) software. The cerebellar electrode was used as a reference, parietal and frontal channels were selected for evaluation.

One week before the experiments, animals acclimatized to the handling procedures and recording setup and mock dosing procedure on four consecutive days. Drug applications started one week after the habituation and a minimum of 3 weeks after the surgery.

### 2.4. Experimental design

We used data from two groups of Lister Hooded rats (from the same supplier, WOBE/Envigo, 8-9 week-old). In the first group, we recorded 12 rats (R1-12, Group_1), but we excluded one rat (R4) from the analysis because of small amplitude EEG. One year later, we measured the second group of Lister Hooded rats (n=6, R1-6, Group_2) and, as we found similar SWDs in their EEG, we dosed them with saline and different drugs to pharmacologically characterize their SWDs. Our recording system is capable of recording six rats simultaneously, so we divided Group_1 into two subgroups and measured them on separate weeks. We measured all the six rats in Group_2 continuously for four weeks. In all measurements, the week started with a habituation day. On this day animals were recorded (data not used), but they received no drug or saline treatments. Both measurements lasted for four experimental weeks. One experimental week contained two (Group_1) or three (Group_2) recording days. In Group_1 measurement a habituation day without any treatment, a control day, when all animal received physiological saline i.p. (0.1 ml/kg), and the third day when all animal received the actual drug (i.p., dissolved or suspended in saline, 0.1 ml/kg). We dosed the different drugs on separate weeks and used the following compounds: *R,S*-baclofen 5 mg/kg, SCH50911 10 mg/kg, diazepam 5 mg/kg, valproate 200 mg/kg (Table 1).

**Table 1.** Experimental design, hab-habituation – no injection only recordings (data not used).

| Week 1 |  | Week 2 |  | Week 3 |  | Week 4 |  |
| --- | --- | --- | --- | --- | --- | --- | --- |
| Mon | Tue | Mon | Tue | Mon | Tue | Mon | Tue |
| habituation | R1-6 | habituation | R7-12 | habituation | R1-6 | habituation | R7-12 |
|  | saline |  | saline |  | saline |  | saline |

**2018 Group\_2**
| Week 1 |  |  | Week 2 |  |  | Week 3 |  |  | Week 4 |  |  |
| --- | --- | --- | --- | --- | --- | --- | --- | --- | --- | --- | --- |
| Mon | Tue | Wed | Mon | Tue | Wed | Mon | Tue | Wed | Mon | Tue | Wed |
| hab | R1-6 | R1-6 | hab | R1-6 | R1-6 | hab | R1-6 | R1-6 | hab | R1-6 | R1-6 |
|  | saline | Baclofen |  | saline | SCH50911 |  | saline | Diazepam |  | saline | Valproate |
|  |  | 5 mg/kg |  |  | 10mg/kg |  |  | 5mg/kg |  |  | 200 mg/kg |

### 2.5. Data analysis

#### 2.5.1. Automatic detection of Spike and Wave discharges (SWDs)

Spike and wave discharges were detected in the EEG of Lister Hooded rats by a MATLAB script based on banded spectral power between 7-15 Hz and 15-30 Hz of the filtered (1-100 Hz) frontal EEG channel. Thresholds for SWD detection were: 3.5 fold of the mean 7-15 Hz power and 3.5 fold of the mean 15-30 Hz power calculated in each rat and only oscillations with duration longer than 1 s were accepted as SWDs (Supplementary Fig 1). The detected SWDs were checked manually on one measurement day in each rat, and the results from automatic detection fitted perfectly with the manual evaluation. Oscillations were counted and classified by a script written in MATLAB.

#### 2.5.2. Characterization of SWDs

Individual SWDs were separated from the EEG of each rat and then their number, duration and average duration were calculated. SWD mean frequency was calculated with MATLAB built-in pwhelch function (hanning window, 2048 frequencies) applied on each SWD followed by detection of the mean peak in the spectrum and then averaged within each animal. The amplitude of individual SWDs was calculated from each SWD, as the difference of the maximal and minimal voltage values of each SWD and averaged afterwards. After drug treatments, changes of SWD parameters were calculated as the difference from control values (measured on the preceding saline day) of each animal.

#### 2.5.3. Sleep stages and EEG power

Sleep stages were classified by a script written in the CED Spike2 software (On/Offline sleep-scoring (*OSD4);* http://ced.co.uk/downloads/scriptspkanal). Sleep stage definitions were adjusted based on parietal EEG and EMG of each rat. The script used band filtered power spectra of the EEG (theta 4.5-9 Hz, delta 2-4.5 Hz) together with the smoothed EMG amplitude. Our 4-hour-long recording period was divided into 20-second-long epochs and categorized into four pre-defined stages. At first, the parietal EEG channel was band-pass filtered (1-100 Hz) and the EMG channel was high pass filtered at 60 Hz. Epochs of wakefulness (WAKE) were defined by high EMG and desynchronized EEG activity. Slow-wave sleep (SWS) was defined by high amplitude delta (2-4 Hz) waves in the EEG and low EMG activity. Rapid eye movement sleep (REM) was defined by very low EMG activity and continuous theta activity in the EEG. Epochs which couldn’t have been classified into any of these 3 stages were defined as DOUBT (mainly showed transitions between stages).

Further data analysis was performed in MATLAB with custom-written scripts. Epochs were counted and normalized to the length of the full recording period in each animal (% time), then grand averages and statistics were calculated.

Power spectral density was calculated from the parietal EEG by MATLAB built-in pwhelch function (hanning window, 2048 frequencies) on each 20-second-long epoch. Individual PSDs to corresponding sleep stages were normalized to the total power in each animal (relative power). Relative power values were averaged first across the 4-hour-long period in each animal. Then, means and standard errors were calculated. Power values in 5, i.e. delta (2-4 Hz), theta (4-9 Hz), alpha (9-15 Hz) and beta (15-30 Hz) and gamma (40-80 Hz) EEG frequency bands were characterized. It is of note that our threshold for evaluation of the lowest frequency range was 2 Hz, thus the 0.5-2 Hz part of the delta frequency band was not quantified in this study.

#### 2.5.4. Statistics

When compared the two different groups of Lister rats, control values were calculated after averaging values measured on two (Group_1) or four (Group_2) days, and then one-way ANOVA, followed by a *post hoc* Tukey-test (in MATLAB) was used for statistical evaluation and an alpha level of 0.05 was set as the significance threshold. In our pharmacological experiments on Group_2, the application of drugs was separated by 1 or 2 weeks and control treatments (saline) were made before each drug application. Thus we assume, that the results could be considered as independent effects. To decrease individual variability we calculated changes from corresponding control values in each animal. In these cases, we used unpaired T-test with test mean of 0 (no change). In the case of sleep stages the values represent % of control so T-test was applied with a test mean of 100.

## 3. Results

### 3.1. Spike-and-Wave discharges

Our automatic SWD detection algorithm based on thresholds in specific frequency bands (3.5 fold as mean level in each rat, both 4-9 Hz and 9-15 Hz bands) detected SWDs in the EEG of each measured Lister Hooded rat (Fig. 1B-E).No significant differences in their numbers or sum and average duration were found between the two groups (Fig 2, SWD number (Fig. 2A): 34-112 / 4 hour in Group_1 and 34-142 /4 hour in Group_2, sum duration (Fig. 2B) 55-298 s/Group_1, 89-186 s/Group_2, average duration (Fig 2C, 1.34-3.52 s / Group_1 and 1.47-1.93s /Group_2). The amplitudes of the discharges were significantly smaller but the peak frequency was higher in Group_2 than in Group_1 (Fig. 2D,E., amplitude: 547 vs 459 μV, p=0.0012, F= 15.79, df: 16, frequency: 7.3 vs 7.9 Hz, p=0.0033, F=12.17, df=16). SWDs were dominant on the frontal and parietal EEG channel of each animal (Fig 1A-E.). The SWD number and duration remained stable for four weeks. (Group_2, Fig 3).

**Fig 1.**
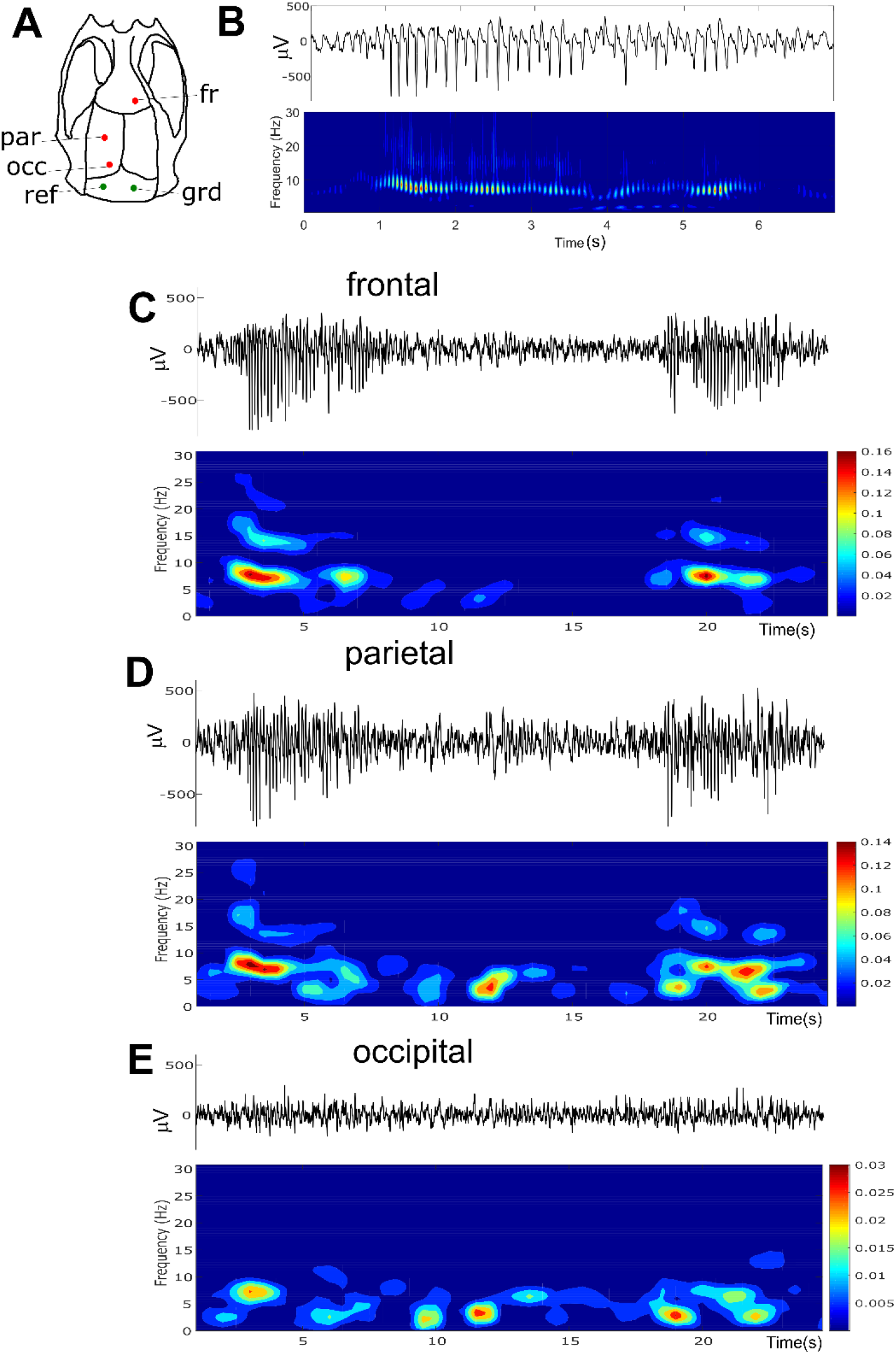
Representative EEG with SWDs measured on three different EEG channels of a Lister Hooded rat. A: electrode placement, B: representative SWD on the frontal electrode and its Morlet wavelet, C – SWDs and their power spectra on the frontal, D - parietal, E – occipital electrodes. Fr-frontal, par-parietal, occ-occipital, grd-ground, ref-reference

**Fig 2.**
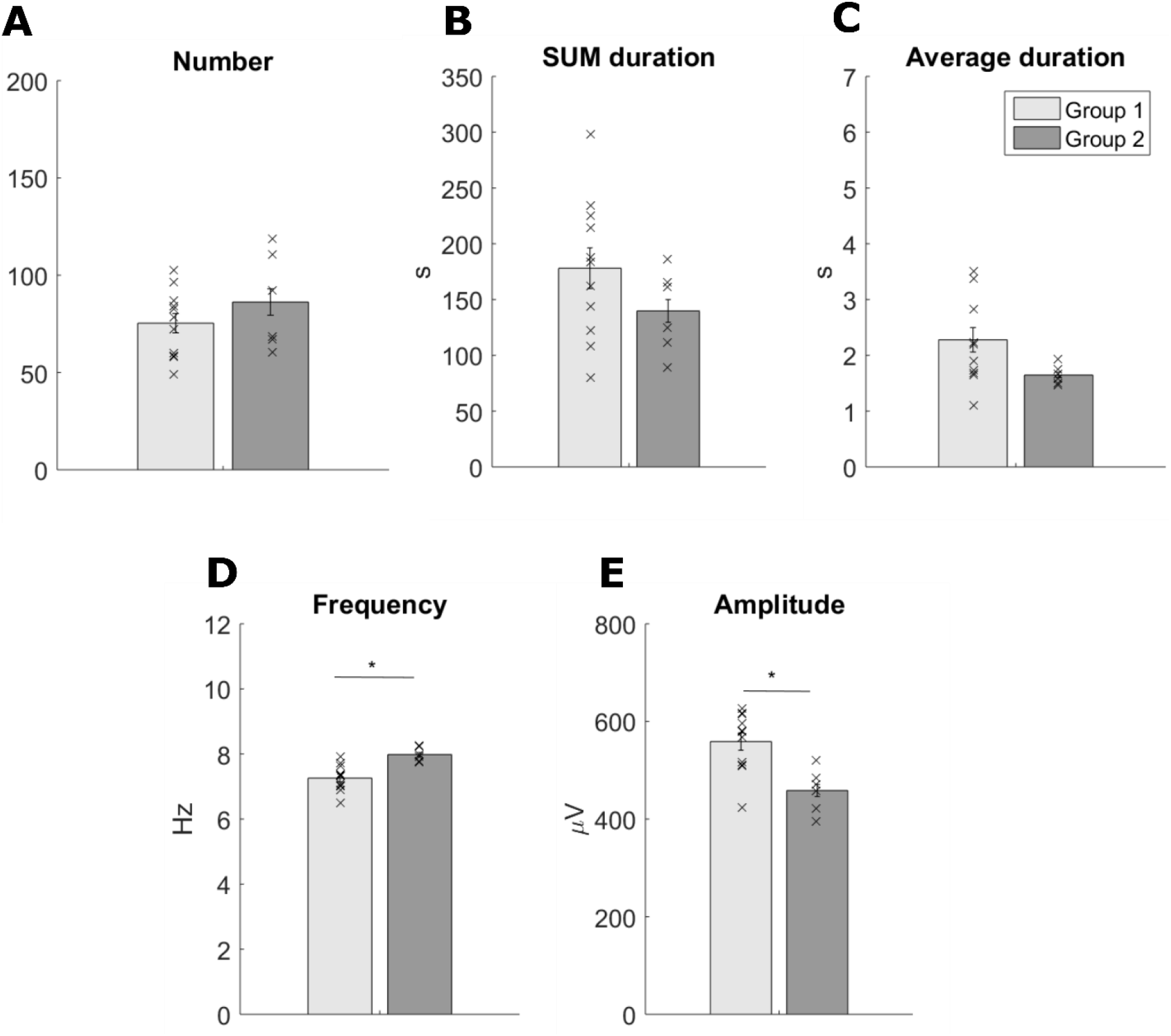
Comparison of different SWD parameters of the two Lister Hooded rat groups. **A) *Number***: all SWD (duration>1 s) counted during the four-hour long recording period. **B) *SUM duration:*** the summed duration of all SWD during the whole recording period. **C) *Average duration***: duration of individual SWDs averaged in each rat. **D)***Frequency*: the frequency peak calculated from the power spectra of each individual SWD and averaged in each rat, then across the rats. **E) *Amplitude***: EEG amplitude measured during SWDs and averaged in each rat. **Group_1:** control data measured 05/02/2017-05/18/2017, n=11, **Group_2**: control data measured on 05/23/2018-06/26/2018, n=6. *-p<0.05, (ANOVA, one-way).

**Fig 3.**
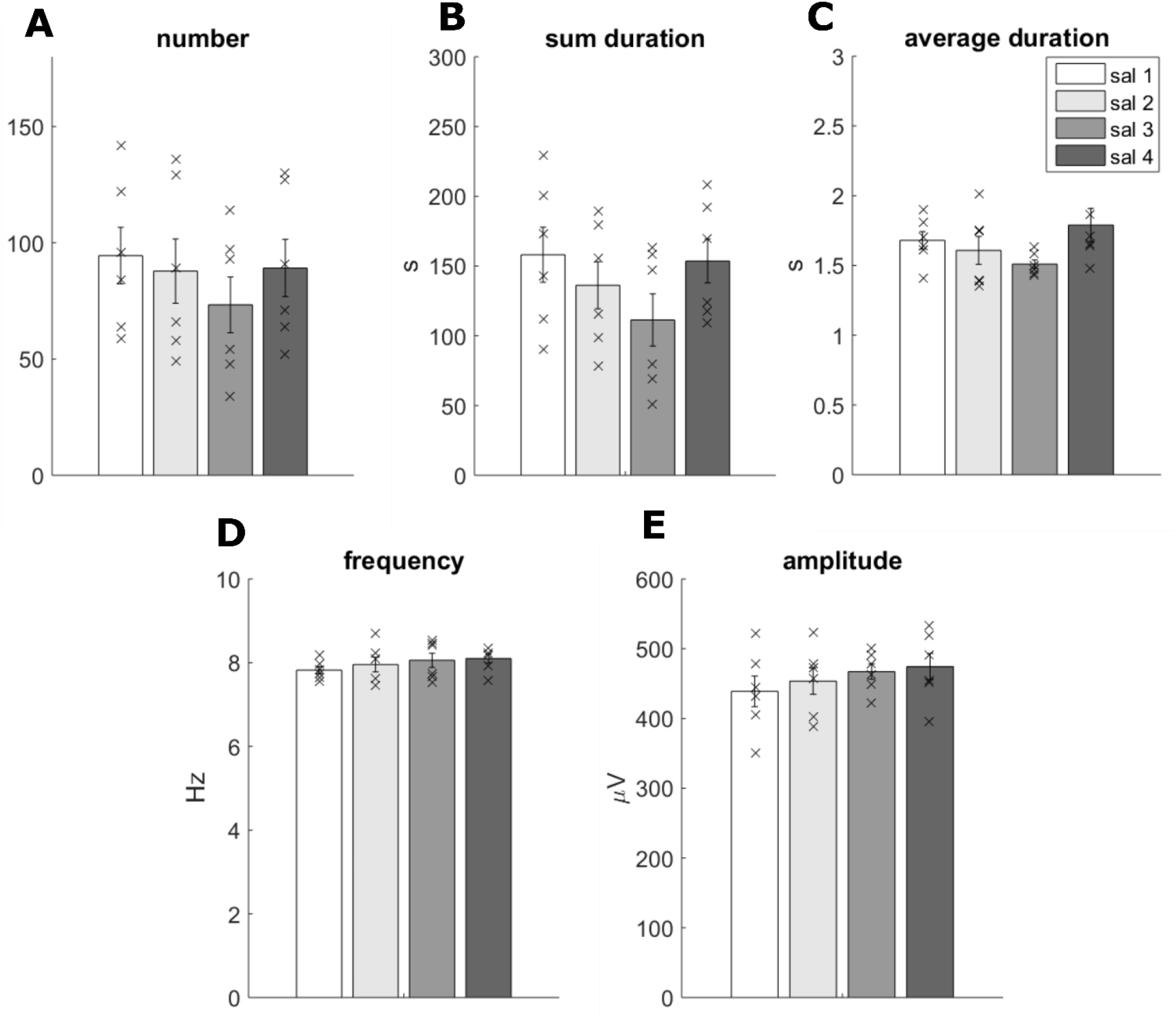
SWD parameters of Group_2 on control days through the 4-week long experiment. **A) *Number***: all SWDs (>1 s-long) counted during the four-hour long recording period. B) ***SUM duration:*** the summarized duration of all SWDs during the whole recording period. C) ***Average duration***: duration of individual SWDs averaged in each rat. D) ***Amplitude***: EEG amplitude measured during SWDs and averaged in each rat. E) *Main frequency*: the frequency peak calculated from the power spectra of each individual SWD and averaged in each rat. Control days were separated by at least one week.

We also grouped the SWDs according to sleep stages during the whole experiment of Group_2 (Table 2,3). The number of SWDs measured in different sleep stages did not differ significantly on control days 1-3 but was higher during REM sleep vs WAKE or SWS on control day 4 (saline-treated days separated by a week, Table 2,3). The sum duration of SWDs was longer during REM than during WAKE or SWS on the 2^nd^ and 4^th^ saline day. The average duration of SWDs was longer during WAKE than during SWS on each control (saline-treated) day. It was longer during WAKE than during REM on the 1^st^ and 4^th^ days (Table 2,3). Interestingly, the main frequency of SWDs was higher during REM than during SWS on days 1 and 2 (Table 2,3).

**Table 2.** Distribution of SWDs (number, sum duration, duration, main frequency) during different sleep stages of Lister Hooded rats. SWD parameters were calculated from control experiments (i.p. saline) made on different weeks during the entire four-hour long recording period.

| **SWD Number** | **Saline 1** |  | **Saline 2** |  | **Saline 3** |  | **Saline 4** |  |
| --- | --- | --- | --- | --- | --- | --- | --- | --- |
| **Sleep stage** | mean | SE | mean | SE | mean | SE | mean | SE |
| **WAKE** | 23.5 | 3.6 | 14.0 | 1.4 | 13.5 | 1.8 | 17.2 | 2.5 |
| **SWS** | 32.8 | 6.6 | 28.8 | 11.1 | 22.8 | 6.0 | 23.5 | 4.8 |
| **REM** | 31.7 | 5.8 | 40.0 | 4.3 | 33.5 | 5.9 | 44.0 | 8.9 |
| **SWD Sum duration (s)** | mean | SE | mean | SE | mean | SE | mean | SE |
| **WAKE** | 52 | 9 | 27 | 4 | 25 | 3 | 46 | 8 |
| **SWS** | 41 | 9 | 36 | 14 | 26 | 7 | 27 | 5 |
| **REM** | 56 | 11 | 67 | 8 | 54 | 12 | 72 | 15 |
| **SWD duration (s)** | mean | SE | mean | SE | mean | SE | mean | SE |
| **WAKE** | 2.2 | 0.16 | 1.9 | 0.23 | 1.9 | 0.22 | 2.6 | 0.16 |
| **SWS** | 1.2 | 0.07 | 1.3 | 0.13 | 1.2 | 0.07 | 1.2 | 0.10 |
| **REM** | 1.8 | 0.12 | 1.7 | 0.05 | 1.5 | 0.10 | 1.6 | 0.08 |
| **SWD frequency (Hz)** | mean | SE | mean | SE | mean | SE | mean | SE |
| **WAKE** | 7.6 | 0.2 | 8.2 | 0.2 | 7.7 | 0.2 | 7.7 | 0.1 |
| **SWS** | 7.4 | 0.2 | 7.2 | 0.4 | 7.9 | 0.3 | 7.8 | 0.3 |
| **REM** | 8.3 | 0.1 | 8.2 | 0.2 | 8.4 | 0.3 | 8.4 | 0.1 |

**Table 3.**
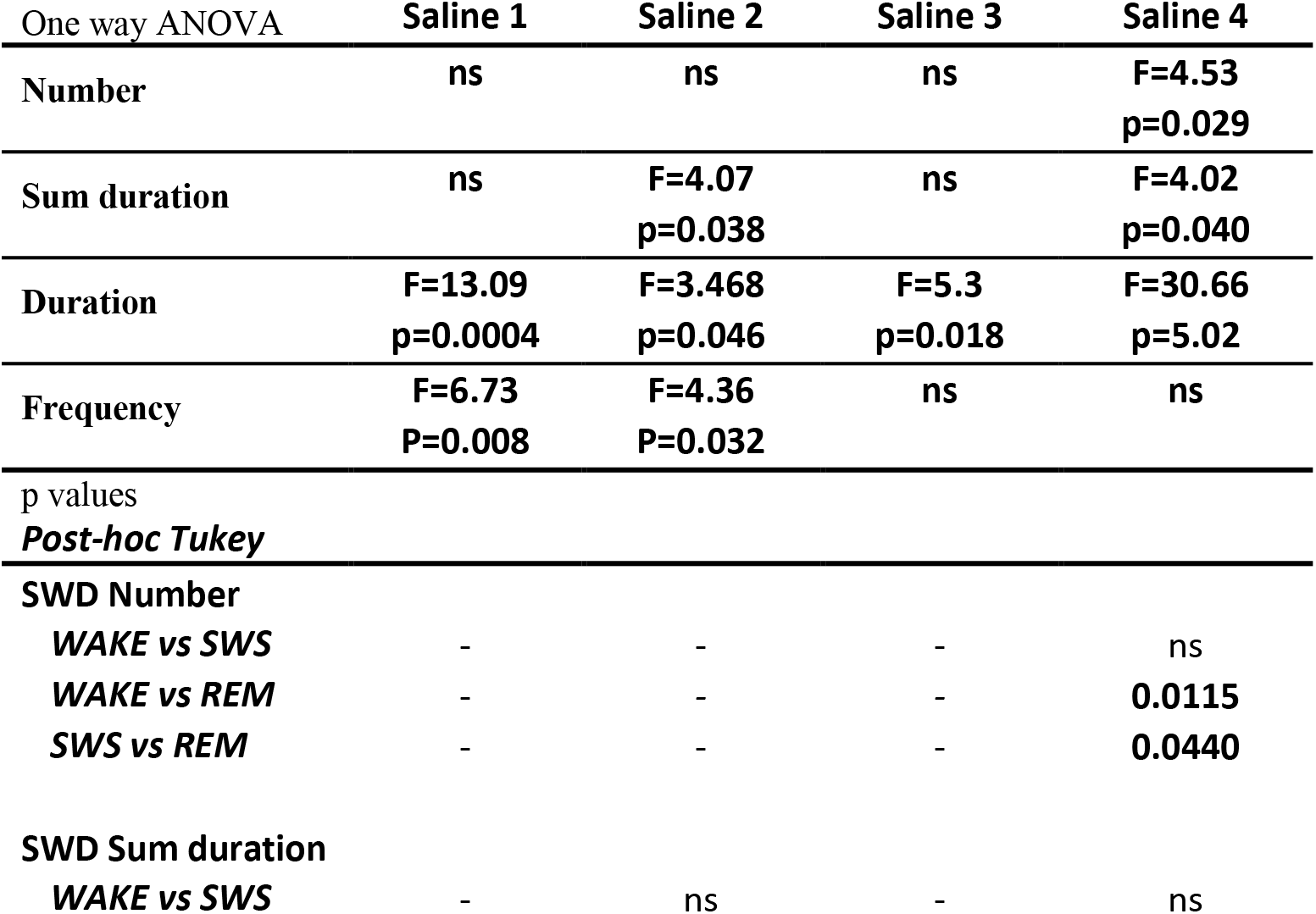
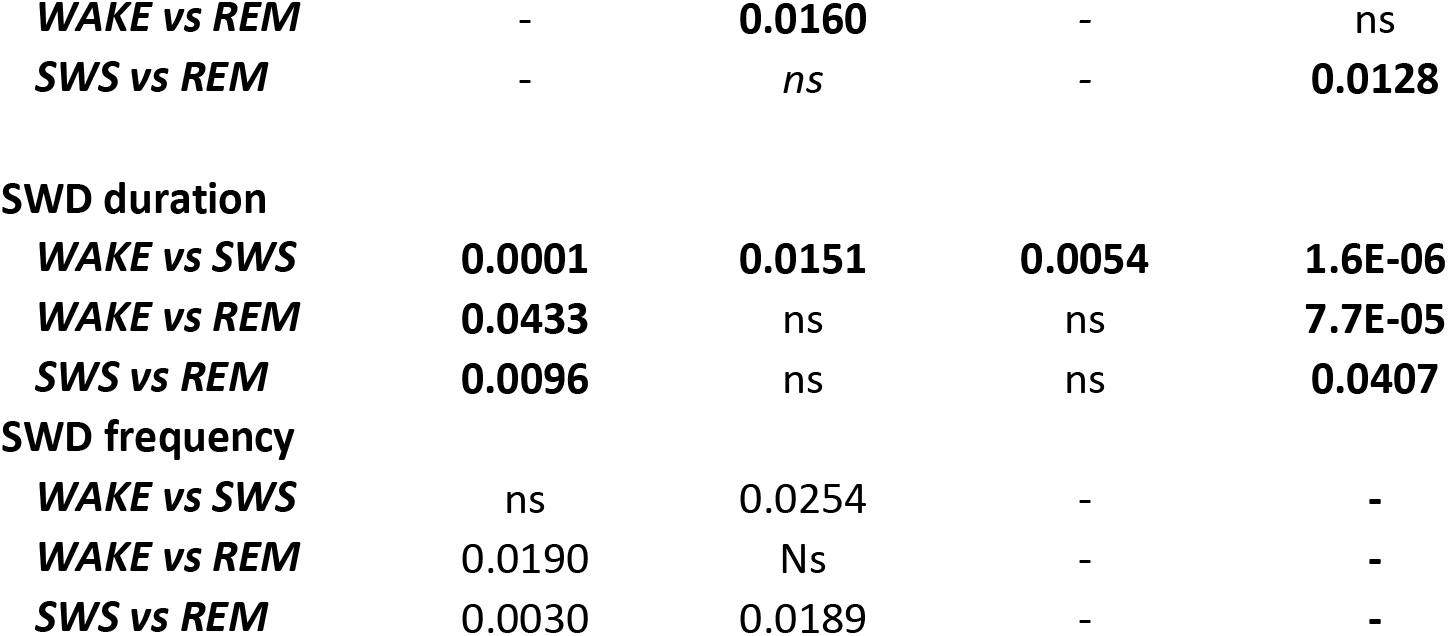
Changes of SWD parameters (number, sum duration, duration, main frequency) during different sleep stages of Lister Hooded rats. SWD parameters were calculated from control experiments (i.p. saline) made on different weeks during the entire four-hour long recording period.

### 3.2. Pharmacological modulation of SWD parameters

#### 3.2.1. GABA-B receptors

First, we calculated the effect of the drugs for each hour during the 4-hour-long recording period to evaluate the temporal characteristics of the effects.

Modulation of the GABA-B receptors had a significant effect on SWD parameters. The GABA-B receptor agonist *R,S*-baclofen (5 mg/kg, i.p.) markedly increased the number and sum duration of SWDs (Hour 1-3, Fig. 4A, B; Supplementary Table 1). The GABA-B receptor antagonist (SCH50911, 10 mg/kg) decreased the number and the sum duration of SWDs (2^nd^ and 4^th^ hour, Fig. 4A, B; Supplementary Table 1). The average duration of SWDs decreased after both compounds (1^st^ and 3^rd^-hour baclofen, 1^st^-3^rd^ hours SCH50911, Fig 2D). SCH50911 also increased the main frequency of SWDs (1^st^, 2^nd^ and 4^th^ hours, Fig 3C) and decreased their amplitude (Supplementary Table 1).

**Fig.4.**
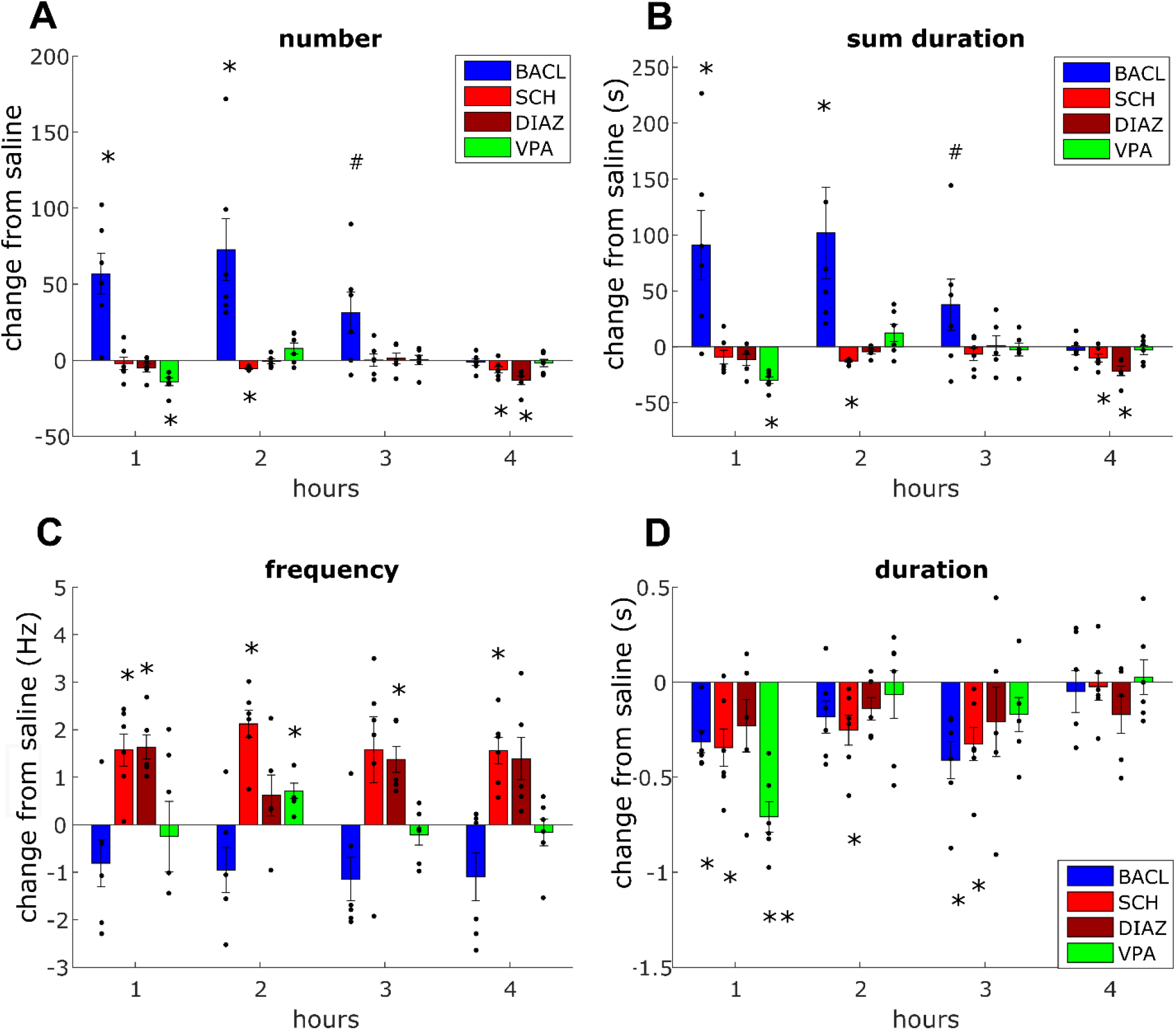
Changes of SWD parameters after application of different drugs. **A:** change in the number of SWDs **B:** change in the sum duration of SWDs **C:** change in the peak frequency, **D:** change in the average duration of SWDs as compared to control (saline) during the 4-hour long recording period. Values depict Mean±S.E.M difference from control (saline) (saline - drug). Colors indicate different drugs applied *i.p.,* blue: *R,S*-baclofen 5 mg/kg, red: SCH50911 10 mg/kg, brown: Diazepam 5mg/kg, green: valproate 200 mg/kg. n=6, * p<0.05, , **p<0.001, # 0.05< p<0.08, T-test.

#### 3.2.2. Anti-epileptic compounds

The anti-epileptic compound, valproate (200 mg/kg) decreased the number and sum duration of SWDs in the first hour (Fig. 4A, B; Supplementary Table 1). It also significantly reduced the average duration of SWDs in the 1^st^ hour (Fig. 4D). Interestingly, valproate slightly increased the peak frequency of SWDs (only in the 2^nd^ hour, Fig. 4C). The GABA-A receptor modulator anti-epileptic drug, diazepam (5 mg/kg) decreased the number and sum duration of SWDs only in the 4^th^ hour while it increased the SWDs peak frequency (in the 1^st^ and 3^rd^ hours, Fig. 4A-C; Supplementary Table 1). SCH50911 decreased the amplitude of SWDs during the whole experiment (Supplementary Table 3).

Temporal changes revealed that the main effects were present 0-2 hours after the drug applications, so we separated the effects according to the sleep stages during the first 2 hours (Fig 5, Supplementary Table 2).

**Fig 5.**
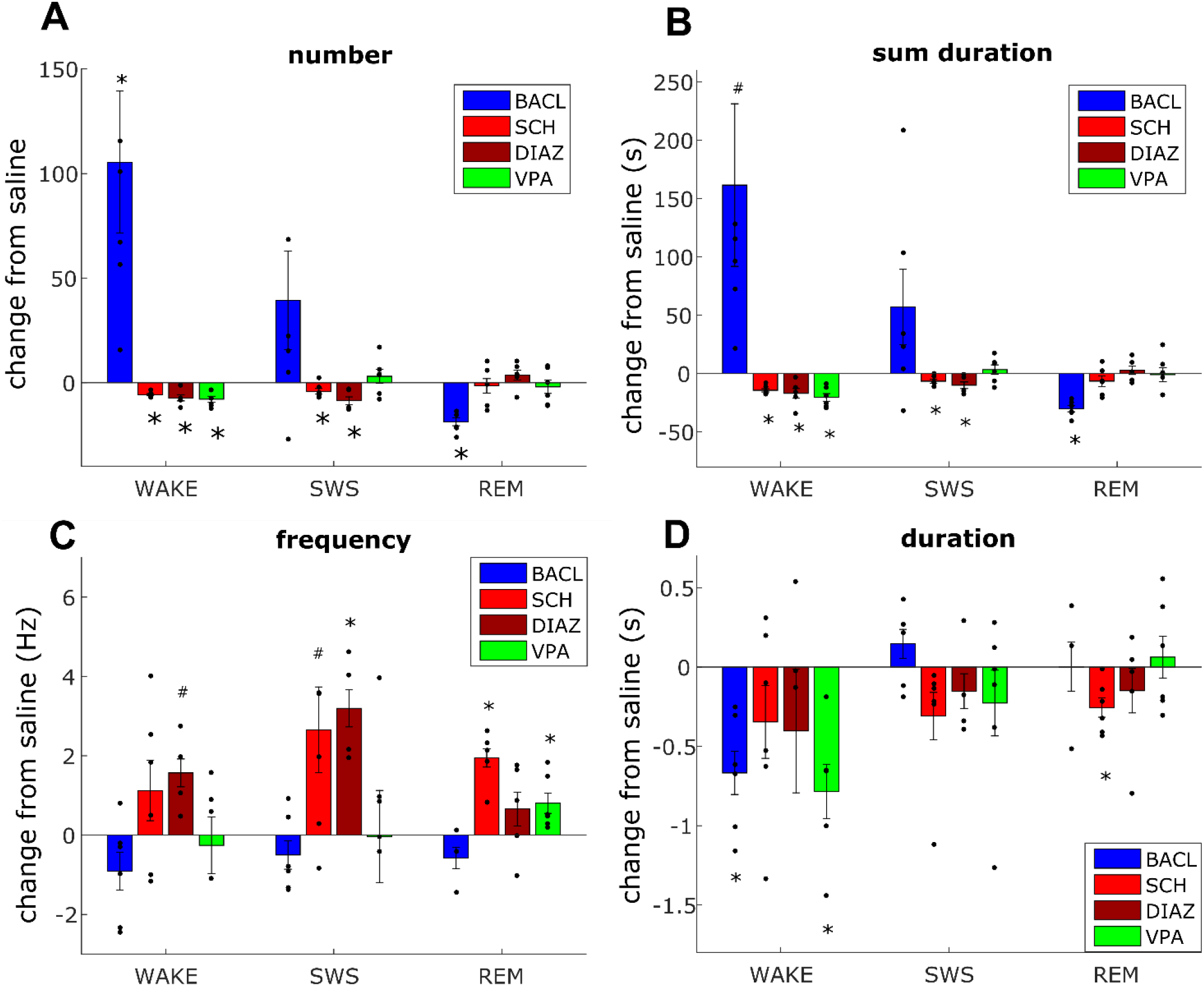
Changes of SWD parameters (from saline) during different sleep stages after application of different drugs. **A:** change in the number of SWDs **B:** change in the sum duration of SWDs **C:** change in the peak frequency, **D:** change in the average duration of SWDs as compared to control (saline) separated according to different sleep stages and measured during the first 2-hour of the recording period. Values depict Mean±S.E.M difference from control (saline, saline-drug). Colors indicate different drugs applied *i.p.,* blue: *R,S*-baclofen 5 mg/kg, red: SCH50911 10 mg/kg, brown: Diazepam 5mg/kg, green: valproate 200 mg/kg. n=6, * p<0.05, **p<0.001, # 0.05< p<0.08, T-test.

This analysis revealed that the enhancement of SWD number by baclofen was predominant in the wake periods (Fig. 5A, B; Supplementary Table 2). Baclofen decreased SWD number and sum duration during REM, but it drastically reduced the time spent in REM sleep too (Table 3, Fig 5A, B; Table 4, Supplementary Table 2).

**Table 4.**
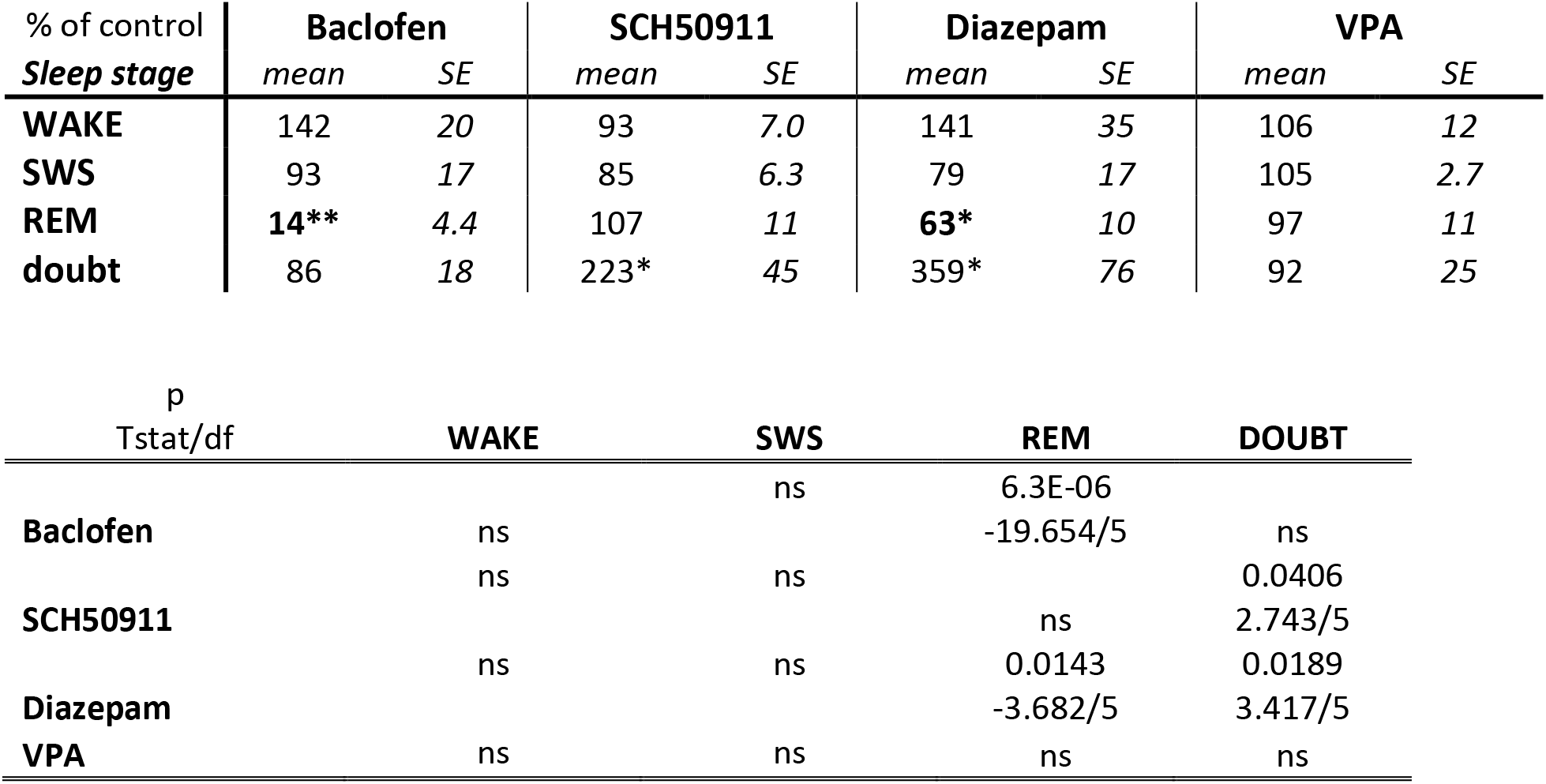
Effects of different drugs on sleep stages of Lister Hooded rats. Time spent in the three different sleep stages during the full 4 hour-long recording period (in percent of values from control recordings). Drugs were applied i.p. before the start of the measurement. Doses were: *R,S*-baclofen 5 mg/kg, SCH50911 10 mg/kg, Diazepam 5 mg/kg, valproate 200 mg/kg. * p<0.05**p<0.001, T-test, test mean: 100.

The anti-epileptic effects became more visible in this analysis: SCH50911 reduced the SWD number and sum duration during both wake and SWS stages and significantly reduced the SWD duration and increased the SWDs peak frequency in REM sleep (Fig. 5A, B; Supplementary Table 2). It also slightly decreased the amplitude of SWDs during REM (Supplementary Table 4). Similarly, diazepam and valproate decreased the number of SWDs during both wake and SWS stages. Diazepam also decreased the sum SWD duration in SWS (Fig 5A, B; Supplementary Table 2), and it also decreased the amplitude of the SWDs during SWS (Supplementary Table 3). Valproate decreased the duration of SWDs during wake (Fig 5D). The peak frequency of SWDs increased after diazepam (SWS) and valproate (REM, Fig. 5C, Supplementary Table 2).

Number and sum duration of SWDs recorded during REM were not affected by SCH50911, valproate or diazepam (Fig 5A, B). But SCH50911 increased the peak frequency and decreased the average duration of SWDs measured during REM (Fig. 5C, D; Supplementary Table 2).

### 3.3. Drug-induced changes in sleep stages

Baclofen and diazepam significantly decreased the time of REM sleep (Table 4). In addition, baclofen and SCH50911 significantly increased the number of doubt epochs (Table 4).

Drugs affected the relative parietal EEG spectral power differently (Supplementary Table 5), baclofen increased the delta (2-4 Hz) and decreased the alpha (9-15 Hz) oscillations, while diazepam and SCH50911 decreased the delta (2-4 Hz) power and increased the relative power of faster frequency bands (beta (15-30 Hz) and gamma (40-80 Hz). SCH50911 additionally increased the alpha power (9-15 Hz, Supplementary Table 5).

## 4. Discussion

In this study, we observed spontaneous SWDs in the EEG of each recorded Lister Hooded rat. The number and duration of the discharges were comparable to that were found in Long Evans or Fischer 344 rats (Ritchie et al., 2003; Shaw, 2004) but were less than reported in inbred genetic models of WAG/Rij and GAERS rat strains (Akman et al., 2010; Drinkenburg et al., 1991).

Long Evans rats were developed by Dr Long through crossing Wistar Institute white female rats with a wild-caught Brown Norway male rat in 1915 (Lindsey and Baker, 2005). Outbred Wistar and Brown Norway rats generate spontaneous SWDs (Willoughby and Mackenzie, 1992) and even wild-caught Brown Norway rats generate them (Taylor et al., 2019). Interestingly, the F1 and F2 hybrids of Fischer 344 and Brown Norway strains generated significantly more and longer duration SWDs than the parental strains (Jando et al., 1995).

The exact origin of the Lister Hooded rat is obscure but maintained at the Lister Institute in the 1920s. It not supposed to be closely related to Long Evans rat. Therefore, the existence of SWD oscillations in the EEG recordings of Lister Hooded rats revealed here is an important finding. At least they are not differing in this sense markedly from Long Evans rats. SWDs can be found up to 90% of Long Evans rats (Shaw, 2004) while we found SWDs in each of the 17 Lister Hooded rats recorded. The average duration of the SWD episodes was around 4 s in Long Evans rats (Shaw, 2004) while 2 s in Lister Hooded rats.

SWDs are spontaneous rhythmic discharges generated by cortico-thalamic circuitry (Blumenfeld, 2003; Steriade, 2005) initiated by the somatosensory cortex (Depaulis and Charpier, 2018; Jarre et al., 2017; Polack et al., 2007; van Luijtelaar et al., 2011). Morphology, distribution, frequency, average duration and sum duration of the discharges we observed were similar to absence epileptic oscillations reported in other strains (Akman et al., 2010; Buzsaki et al., 1988a; Drinkenburg et al., 1991; Jando et al., 1995; Ritchie et al., 2003; Rodgers et al., 2015; Shaw, 2004; Willoughby and Mackenzie, 1992). In our recordings, the discharges were apparent on the frontal electrodes and clearly visible in the parietal areas but smaller over the occipital cortex as reported in case of typical absence oscillations in rats (Blumenfeld, 2003; Meeren et al., 2002).

Although SWD related 7–12 Hz oscillations in Long Evans rats have been observed repeatedly for many decades, their role remains a matter of debate. An alternative interpretation of the role of these oscillations during quiet immobility suggests that it would be a dynamic filter optimized for detecting weak or novel tactile stimuli from vibrissae (Fanselow et al., 2001; Nicolelis et al., 1995; Wiest and Nicolelis, 2003). This hypothesis proposes that under an awake behavioural state, they are a functional analogue of the physiological mu rhythm of humans that not observed during sleep. Consequently, evaluation of the occurrence of SWDs during sleep states could define their possible functional correlates (Shaw, 2004). In Long Evans rats, SWDs appeared during SWS and REM sleep and in the short-lasting period of vigilance change (Shaw, 2004). SWDs are commonly observed in this transition state of wakefulness to sleep in inbred absence epileptic rats (Danober et al., 1998; Drinkenburg et al., 1991). In our study, SWDs occurred in Lister Hooded rats in each sleep stages, and we did not observe significant stage preference.

Anti-epileptic drugs typically used to verify experimental epileptic animal models, e.g. diazepam and valproate, successfully suppress the occurrence of SWD in human as well as in Long-Evans rats (Shaw, 2007). Similarly, in genetically epileptic animal models (in WAG/Rij and GAERS rats) and absence epileptic patients the GABA-B receptor agonist baclofen augments SWDs while the GABA-B receptor antagonist reduces them (Manning et al., 2003; Marescaux et al., 1992a; Micheletti et al., 1985; Ritchie et al., 2003; Vergnes et al., 1997). In our study, in Lister Hooded rats, the pharmacological treatments show similar results: the GABA-B receptor agonist baclofen augmented the SWDs while the GABA-B receptor antagonist, as well as diazepam and valproate, reduced them.

Nevertheless, when we examined the effects of these pharmacological agents on the detected SWDs according to the sleep stages, we have found some differences. Diazepam and valproate did not affect the SWDs during REM sleep while decreased their number and sum duration during wake and SWS. The SCH50911 altered the SWDs characteristics measured during REM: it increased their peak frequency and decreased their average duration. Valproate selectively increased the frequency of SWDs during REM. Therefore, the SWDs during REM sleep may have distinct neurophysiological characteristics that are different from common absence-like SWDs.

The increased peak frequency of SWDs observed by SCH50911 (SWS, REM), diazepam (wake, SWS) and valproate (REM) might be explained by the modification of the intrinsic bursting properties of the cortico-thalamo-cortical network by GABA-B and GABA-A receptors, in line with previous results (Manning et al., 2003; Marescaux et al., 1985; Marescaux et al., 1992a; Ritchie et al., 2003; Shaw, 2007; Vergnes et al., 1997). The change in the dominant action of GABA-A and GABA-B on thalamo-cortical neurons demonstrated to cause a shift in the oscillation frequency of SWDs in a computational model (Destexhe, 1999).

In our study, diazepam also increased the EEG spectral power in the beta and gamma frequency band and reduced delta power as reported previously (Siok et al., 2012; Tobler et al., 2001; van Lier et al., 2004). This effects of diazepam on the EEG spectral power were similar to those of the GABA-B receptor antagonist SCH50911. Inhibition of the thalamic GABA-B receptors can decrease slow EEG oscillations (Juhasz et al., 1994). Our results also revealed that modulation of the GABA-B receptor altered the power in the alpha frequency band.

We also observed changes in the ratio of sleep stages, in line with previous results: diazepam increased the number of non-classified epochs (doubt) and decreased the amount of REM sleep (Tobler et al., 2001; van Lier et al., 2004), and baclofen decreased the time spent in REM sleep drastically (Ulloor et al., 2004).

Lister Hooded rats predominantly used in preclinical behavioural pharmacology experiments like in touchscreen-based visual discrimination tasks (Mohler et al., 2015; Talpos et al., 2012). The comparisons with the two most widely used albino rat strains revealed differences in tests of general locomotor activity, namely a significantly higher daily activity and lower nocturnal activity for Lister Hooded rats compared to the others (McDermott and Kelly, 2008). A strain comparison with Long Evans rats also conducted in touchscreen-based translational rodent experiments. In this study, Lister Hooded rats were more sensitive than Long Evans rats to three out of four cognition-impairing drugs tested (Mohler et al., 2015). Our results might explain the difficulty some scientists may have in reproducing some findings on different rat strains, especially when the existence of probable SWDs on the cortical EEG is uncertain.

Recording the EEG of the rats during behavioural experiments is not a common approach. Therefore, the existence of the SWDs or their modification by drugs could result in puzzling behavioural responses. Pharmacological agents could cause specific changes in SWD oscillations (Bortolato et al., 2010; Ritchie et al., 2003; Vergnes et al., 1997). During SWDs the functional state of the cerebral cortex should be depressed (Shaw, 2004), in line with their absence epileptic-like nature, as on several occasions, ongoing behaviour such as exploratory sniffing is interrupted during SWDs and then continued after it (Willoughby and Mackenzie, 1992). Some early studies reported preserved learning and social skills in epileptic strains (Vergnes et al., 1991), while currently existence of altered behaviour and impaired cognitive performance is widely accepted in genetically absence epileptic rats (Roebuck et al., 2020; Russo and Citraro, 2018). The cognitive impairment in the WAG/Rij rat strain might be secondary to the occurrence of absence seizures (Leo et al., 2019). In Lister Hooded rats, generally higher activity was described (McDermott and Kelly, 2008). Interestingly, it can manifest in a specific acoustically elicited bursts of locomotor running (Commissaris et al., 2000; Neophytou et al., 2000). This acoustically elicited hyper-responsiveness is quite the opposite of what a brief spontaneous paroxysmal SWD would induce: behavioural hypo-responsiveness. Nevertheless, the described SWDs in the Lister Hooded rat could still mark an “epileptic” abnormal phenomenon as longer duration, higher intensity acoustic stimulus or longer duration white noise induce convulsions in a significant number of animals (Commissaris et al., 2000).

In conclusion, we described SWDs on the EEG of Lister Hooded rats similarly to Long Evans rats that do not explain their strain differences. The SWDs occurred in each sleep stages, and pharmacological agents modulated their occurrence as it had found in absence epileptic rat strains. Although the implication of the SWDs is not clear yet, the modulation of their incidence, in pharmacological behavioural experiments, should also be considered at the evaluation of the drug effects.

## Abbreviations

SWD: spike and wave discharge
SWS: Slow Wave Sleep
REM: Rapid Eye Movement Sleep
GAERS: Genetic Absence Epilepsy Rat of Strasbourg
Wag/Rij: Wistar Albino Glaxo Rats from Rijswijk

## 6. Acknowledgements

The authors are grateful for the technical assistance of Petra Schreiber and Katalin Tóthné Fekete.

Supplementary Information accompanies this paper.

## 7. Author contributions

A.Cz. and G.Ny. designed the experiments; P.D. performed the data acquisition; G.Ny., G.S. and P.D. analysed the data;. A.Cz. and G.Ny wrote the main manuscript text. All authors reviewed the manuscript.

---

**Supplementary Fig 1.**
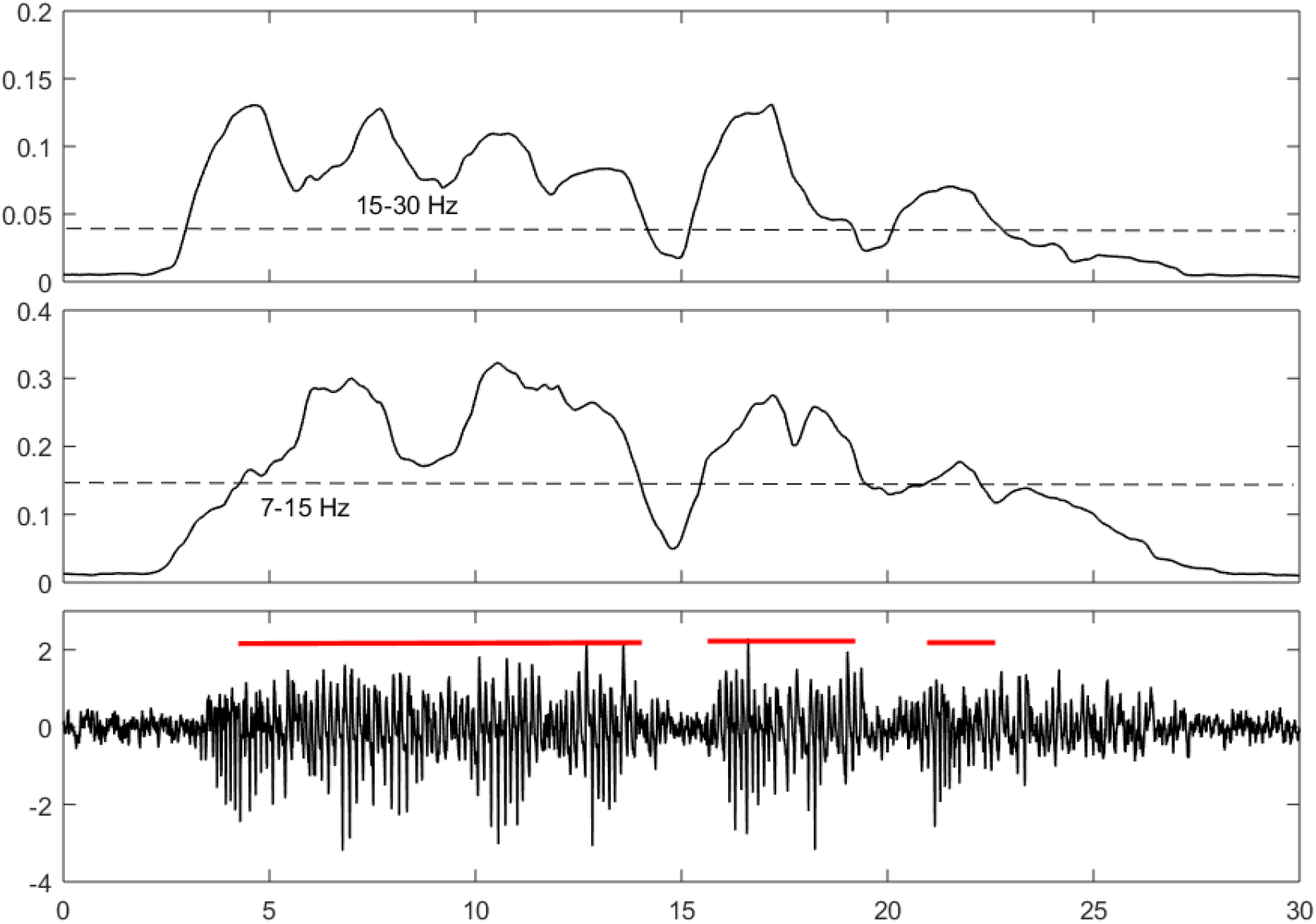
SWD detection based on thresholds set on the banded power values of two frequencies (7-15 and 15-30 Hz). TOP: power of frontal EEG filtered between 7 and 15 Hz. Middle: power of the frontal EEG filtered between 15-30 Hz. Dashed lines depicts the corresponding thresholds (3.5x mean) for SWD detection. Bottom: frontal EEG with SWD s. Red line depicts the detected SWD durations.

**Supplementary Table 1.** Statistical analysis **of SWD parameters after application of different drugs. (Fig 4). T test was applied in each case, test mean: 0, ns: p>0.05.**

| p<br>Tstat/df | hour1 | hour2 | hour3 | hour4 |
| --- | --- | --- | --- | --- |
| **SWD NUMBER** |  |  |  |  |
|  | 0.0115 | 0.0222 |  |  |
| **Baclofen** | 3.894/5 | 3.269/5 | ns | ns |
|  |  | 0.0001 |  | 0.0394 |
| **SCH50911** | ns | 10.51/5 | ns | -2.769/5 |
|  |  |  |  | 0.0179 |
| **Diazepam** | ns | ns | ns | 3.872/4 |
|  | 0.0041 |  |  |  |
| **VPA** | -5.01/5 | ns | ns | ns |
| **SWD Sum Duration** |  |  |  |  |
| **Baclofen** | 0.0436 |  | ns | ns |
|  | 2.684/5 | ns |  |  |
| **SCH50911** |  | 3.57E-05 |  | 0.0478 |
|  | ns | 13.812/5 | ns | 2.609/5 |
| **Diazepam** |  |  |  | 0.0137 |
|  | ns | ns | ns | 4.203/4 |
| **VPA** | 0.0003 | ns | ns | ns |
|  | 9.134/5 |  |  |  |
| **SWD Duration** |  |  |  |  |
| **Baclofen** | 0.0040 |  | 0.0120 | ns |
|  | -5.038/5 | ns | -3.852/5 |  |
|  | 0.0244 | 0.0319 | 0.0191 | ns |
| **SCH50911** | -3.185/5 | -2.95/5 | -3.406/5 |  |
| **Diazepam** | ns | ns | ns | ns |
| **VPA** | 0.0005 | ns | ns | ns |
|  | -8.080/5 |  |  |  |
| **SWD Frequency** |  |  |  |  |
| **Baclofen** | ns | ns | ns | ns |
| **SCH50911** | 0.0082 | 0.0011 |  | 0.0037 |
|  | 4.237/5 | 6.734/5 | ns | 5.13/5 |
| **Diazepam** | 0.0063 |  | 0.0149 | ns |
|  | 5.246/4 | ns | 4.97/4 |  |
| **VPA** |  | 0.0106 |  | ns |
|  | ns | 3.974/5 | ns |  |

**Supplementary Table 2.**
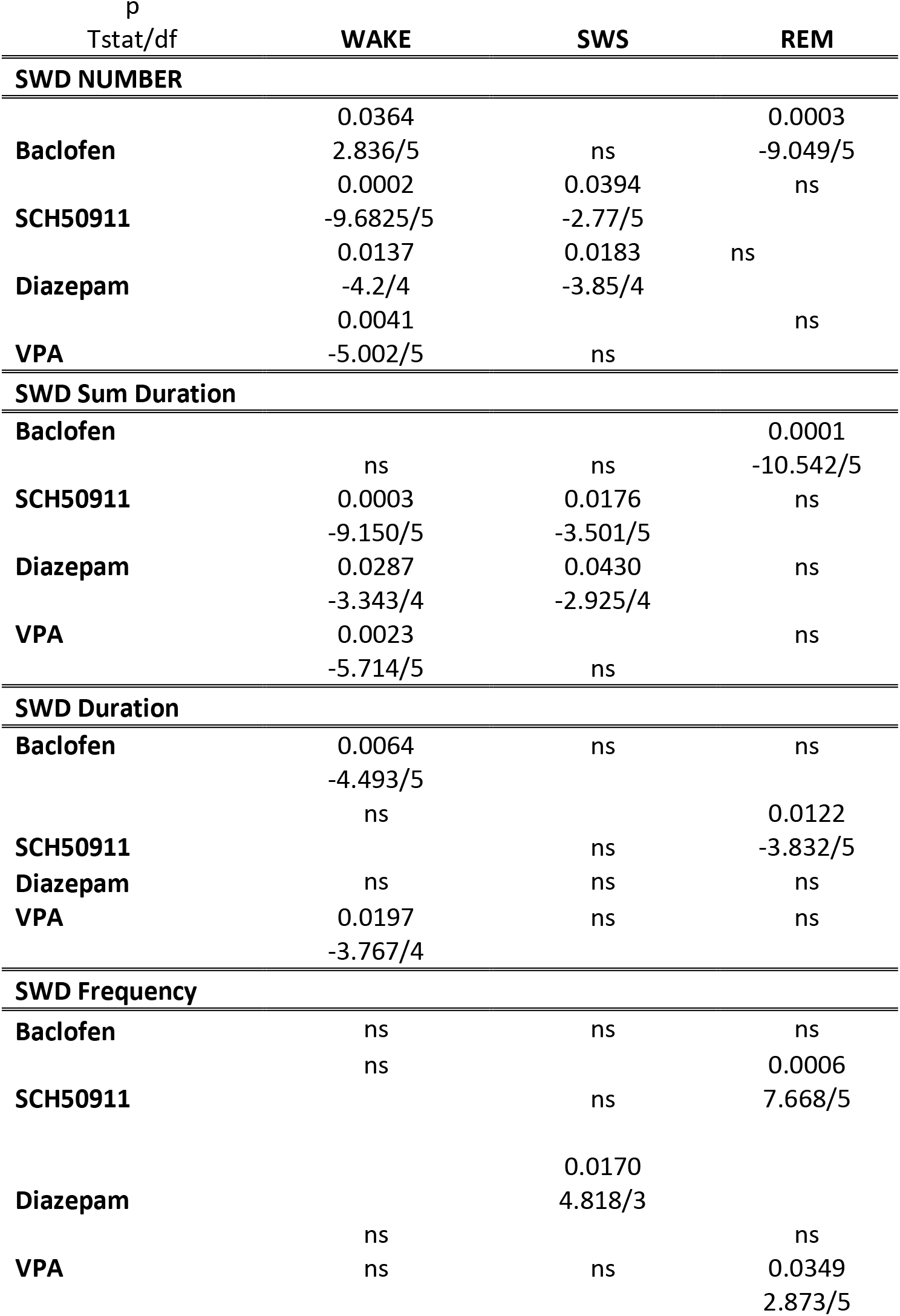
Statistical analysis **of SWD parameters during different sleep stages. (Fig 5). T test was applied in each case, test mean: 0, ns: p>0.05. df: degrees of freedom.**

**Suppl Table 3.**
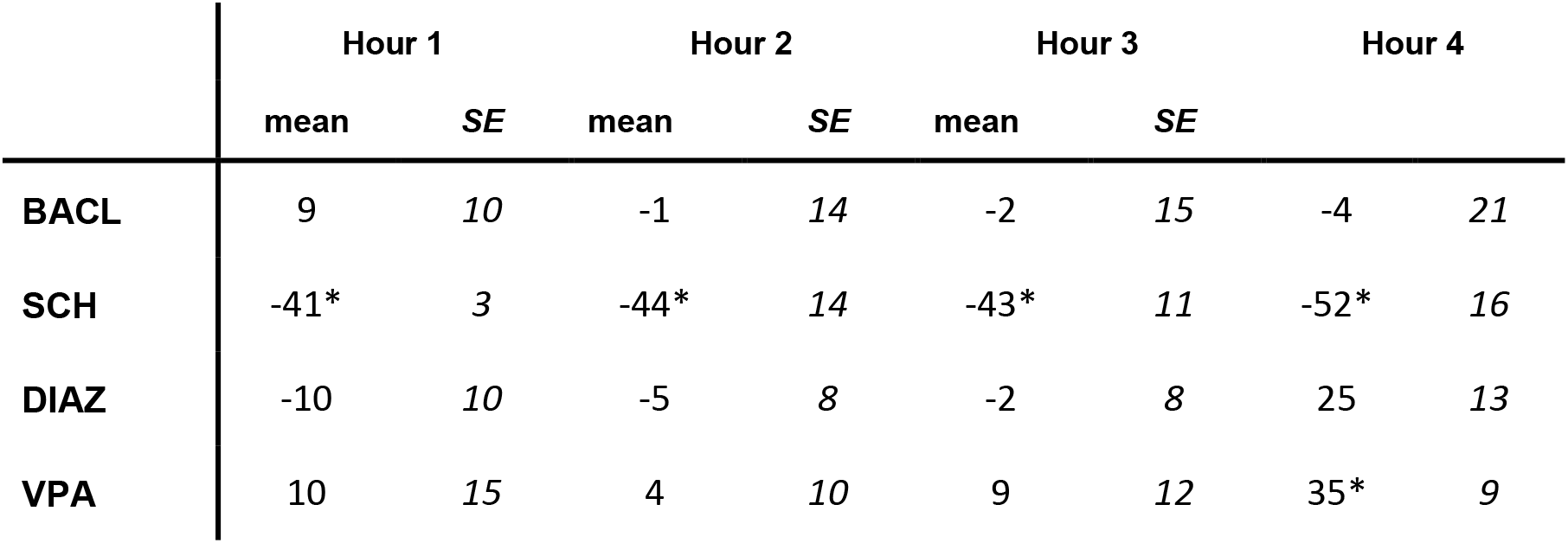
Changes of SWD amplitude (from saline,μV) averaged in each hour after application of different drugs. **T test was applied in each case, test mean: 0, * p<0.05. SCH50911 hour1: p=9.3E-05, tstat=-11.36, hour2: p=0.0339, t=−2.89, hour3: p=0.0137, t=−3.722, hour4: p=0.0327, t=−2.927, hour1-4 degrees of freedom=5. VPA hour 4: p=0.0141, t=3.687, df=5.**

**Suppl Table 4.**
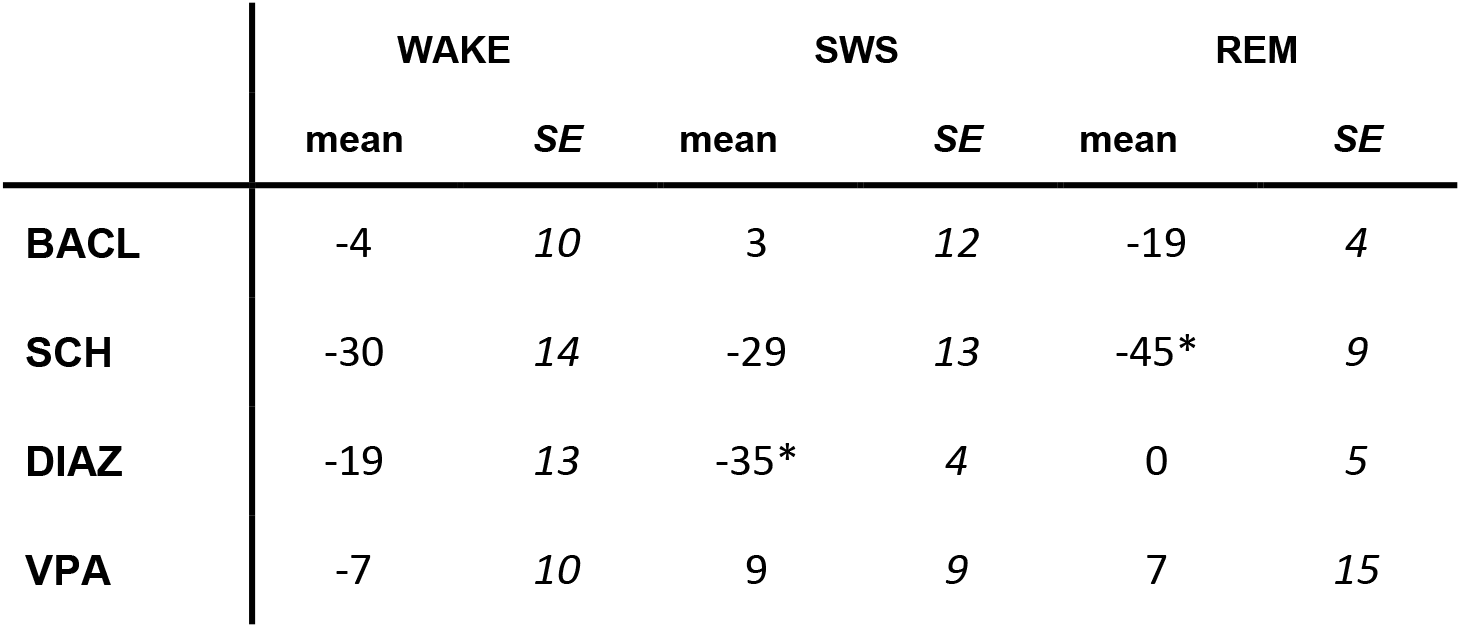
Changes of SWD amplitude (from saline, μV) averaged in different sleep stages after application of different drugs. **T test was applied in each case, test mean: 0, * p<0.05. SCH, REM: p=0.0065, t=−4.484, df=5, Diazepam, SWS: p=0.0094, t=−5.957, df=3.**

|  | WAKE |  | SWS |  | REM |  |
| --- | --- | --- | --- | --- | --- | --- |
|  | mean | SE | mean | SE | mean | SE |
| **BACL** | -4 | 10 | 3 | 12 | -19 | 4 |
| **SCH** | -30 | 14 | -29 | 13 | -45* | 9 |
| **DIAZ** | -19 | 13 | -35* | 4 | 0 | 5 |
| **VPA** | -7 | 10 | 9 | 9 | 7 | 15 |

**Supplementary Table 5.** Changes in EEG power spectral density (normalized) of Lister Hooded rats after different treatments calculated from the whole recording period (4 hours). Power spectral density of the five frequency bands, Delta (2-4 Hz), Theta (4-9 Hz), Alpha (9-15 Hz), Beta (15-30 Hz) and gamma (40-80 Hz) were calculated from the filtered (1-100 Hz) parietal EEG and normalized to all power in each animal. SAL: saline, BACL: *R,S*-baclofen 5 mg/kg, SCH: SCH50911 10 mg/kg, DIAZ: Diazepam 5 mg/kg, VPA: valproate 200 mg/kg. Anova, # −0.05<p<0.09, * −p<0.05, ** −p<0.001.

|  | DELTA |  | THETA |  | ALPHA |  | BETA |  | GAMMA |  |
| --- | --- | --- | --- | --- | --- | --- | --- | --- | --- | --- |
|  | mean | SE | mean | SE | mean | SE | mean | SE | mean | SE |
| **SAL** | 0.2684 | 0.01623 | 0.3955 | 0.01360 | 0.1847 | 0.00636 | 0.1030 | 0.00279 | 0.0323 | 0.00205 |
| **BACL** | **0.3049<sup>#</sup>** | 0.01204 | 0.3838 | 0.00822 | **0.1593\*** | 0.00493 | 0.1061 | 0.00611 | 0.0304 | 0.00251 |
| **SCH** | **0.1913\*\*** | 0.01347 | 0.3963 | 0.01934 | **0.2100\*** | 0.00465 | **0.1279\*** | 0.00547 | **0.0502\*** | 0.00467 |
| **DIAZ** | **0.1900\*\*** | 0.01005 | 0.3740 | 0.02059 | 0.1968 | 0.00570 | **0.1538\*\*** | 0.00769 | **0.0508\*** | 0.00432 |
| **VPA** | 0.2849 | 0.02126 | 0.3764 | 0.01449 | 0.1812 | 0.00705 | 0.1045 | 0.00368 | 0.0355 | 0.00243 |

